# Conserved cysteine residues in Kaposi’s sarcoma herpesvirus ORF34 are necessary for viral production and viral pre-initiation complex formation

**DOI:** 10.1101/2023.03.08.531831

**Authors:** Tadashi Watanabe, Aidan McGraw, Kedhar Narayan, Hasset Tibebe, Kazushi Kuriyama, Mayu Nishimura, Taisuke Izumi, Masahiro Fujimuro, Shinji Ohno

## Abstract

Kaposi’s sarcoma herpesvirus (KSHV) ORF34 plays a significant role as a component of the viral pre-initiation complex (vPIC), which is indispensable for late gene expression across beta and gamma herpesviruses. Although the key role of ORF34 within the vPIC and its function as a hub protein have been recognized, further clarification regarding its specific contribution to vPIC functionality and interactions with other components is required. This study employed a deep-learning algorithm-assisted structural model of ORF34, revealing highly conserved amino acid residues across human beta- and gamma-herpesviruses localized in structured domains. Thus, we engineered ORF34 alanine-scanning mutants by substituting conserved residues with alanine. These mutants were evaluated for their ability to interact with other vPIC factors and restore viral production in cells harboring the ORF34-deficient KSHV-BAC. Our experimental results highlight the crucial role of the 4 cysteine residues conserved in ORF34: a tetrahedral arrangement consisting of a pair of C-X_n_-C consensus motifs. This suggests the potential incorporation of metal cations in interacting with ORF24 and ORF66 vPIC components, facilitating late gene transcription, and promoting overall virus production by capturing metal cations. In summary, our findings underline the essential role of conserved cysteines in KSHV ORF34 for effective vPIC assembly and viral replication, thereby enhancing our understanding of the complex interplay between the vPIC components.

**IMPORTANCE:** The initiation of late gene transcription is universally conserved across the gamma- and beta-herpesvirus families. This process employs a viral pre-initiation complex (vPIC), which is analogous to a cellular PIC. Although KSHV ORF34 is a critical factor for viral replication and is a component of the vPIC, the specifics of vPIC formation and the essential domains crucial for its function remain unclear. Structural predictions suggest that the 4 conserved cysteines (C170, C175, C256, and C259) form a tetrahedron that coordinates the metal cation. We further investigated the role of these conserved amino acids in interactions with other vPIC components, late gene expression, and virus production, to demonstrate for the first time that these cysteines are pivotal for these functions. This discovery not only deepens our comprehensive understanding of ORF34 and vPIC dynamics but also lays the groundwork for more detailed studies on herpesvirus replication mechanisms in future research.

## Introduction

Human herpesviruses are classified into the *Alpha*-, *beta*-, and *Gamma*-*herpesvirinae* subfamilies (1, 2). Kaposi’s sarcoma herpesvirus (KSHV), also referred to as human gammaherpesvirus-8 (HuGHV8) or human herpesvirus-8 (HHV-8), along with Epstein-Barr virus (EBV), belongs to the *Gammaherpesvirinae* subfamily (1–3). Unique among human herpesviruses, these human gamma herpesviruses possess direct pathogenicity toward malignant neoplasms (3–5). KSHV plays a notable role in the pathogenicity of epithelial malignancies with Kaposi’s sarcoma (KS), and of B-lymphocyte malignancies with multicentric Castleman’s disease (MCD) and primary effusion lymphoma (PEL) (4, 6–9). Additionally, KSHV infection can lead to an inflammatory disorder known as KSHV inflammatory cytokine syndrome (KICS) (6–8).

The KSHV genome encodes more than 90 gene products, including proteins and functional RNAs such as lncRNAs, miRNAs, and circRNAs (6, 10–14). The majority of these genes are crucial for the KSHV life cycle, which incorporates 2 distinct phases: latent and lytic. After primary infection, KSHV initially expresses a few genes known as latent genes, which contribute to establishing and maintaining latency within infected cells (15). The transition from a latent to lytic infection state necessitates certain reactivation stimuli to infected cells, such as extracellular environmental changes and intracellular signaling alterations that result in the upregulation of a multitude of KSHV-coding genes, referred to as lytic genes (16–18). In association with the progress of virus-producing infection, the expression pattern of lytic genes is divided into 3 sequential steps, resulting in the classification of lytic genes into immediate-early (IE), early (E), and late (L) genes (6, 19, 20). The initial response to reactivation stimuli is the upregulation of IE-genes, including the viral transcription factor ORF50/K-Rta, which instigates the lytic infection state. ORF50/K-Rta is known to robustly activate various lytic gene promoters, including E-genes (21, 22). The subsequent expression of E-genes leads to replication of the viral DNA genome, as certain E-genes encode viral DNA polymerase factor proteins. This process then paves the way for the transcription of L-genes, encoding various viral structural proteins (*e.g.*, capsid, tegument, and envelope proteins), and ensuring the efficient production of progeny virus particles (6, 23).

A unique regulatory system for the expression of L-genes has been observed in gamma- and beta-herpesviruses, but not in alpha-herpesviruses. Most viruses employ the host transcription machinery, including the pre-initiation complex (PIC) consisting of TATA-binding protein (TBP) bound to specific DNA sequences, and general transcription factors (GTFs), which utilize the host PIC machinery for the expression of IE- and E-genes. Conversely, these 2 herpesvirus subfamilies rely on a viral PIC (vPIC) for the transcription of some L-genes (24–26).

In KSHV, 6 viral proteins (ORF18, ORF24, ORF30, ORF31, ORF34, and ORF66) have been identified as components of vPIC (27–33). KSHV ORF24 is thought to be a functional analog of TBP, since ORF24 directly binds to a TATA-box-like DNA sequence known as the TATT motif, which is present at the transcription start site (TSS) of some L-genes (25). ORF24 recruits the RNA polymerase II (RNAPII) complex to the L-gene TSS and initiates mRNA transcription (28, 34). In contrast, KSHV ORF34 interacts directly with ORF24 and serves as a binding platform for several PIC components, namely ORF18, ORF31, and ORF66 (27–33). ORF30 seems to directly bind only to ORF18, as association with the other vPIC components has not been detected (28, 33, 35). Each component of this protein complex is indispensable for the expression of the L-gene, which is essential for infectious virus production (27–33). However, the precise interaction mechanism among each vPIC component and the stepwise process of assembly into the complete vPIC remain to be elucidated. In addition to recruiting RNAPII to the L-gene TSS, other functions of vPIC are poorly understood.

Our previous study explored the role of KSHV ORF34 in viral replication, revealing it as a binding platform for vPIC components (27). We demonstrated that the central region of ORF34 was critical for binding to ORF18, ORF31, and ORF66. We also found that the C-terminal region of ORF34 was necessary for interaction with ORF24 and virus production. These results indicate that ORF34 assembles several vPIC components (ORF18, ORF31, and ORF66) via its middle region, and its C-terminus harnesses them to ORF24 to form a vPIC at the L-gene TSS. However, detailed information regarding the molecular structure of ORF34 and its contribution to its function remains unknown.

To elucidate how the microstructure and/or domain structure of ORF34 influences its physiological functions, a deep-learning-based structural model of ORF34 was initially generated. This model predicted a tetrahedral arrangement of the 4 conserved cysteines (C170, C175, C256, and C259) in ORF34 that are potentially instrumental in the incorporation of metal cations. Subsequent experimental evaluation of this predicted model demonstrated that these conserved cysteines were instrumental in vPIC assembly, late gene transcription, and viral production.

## Results

### Predicted structure model of KSHV ORF34 utilizing deep-learning algorithm and identification of conserved amino-acid residues between ORF34 and beta- and gamma-herpes viral homologs

Based on our previous study (27), we attempted to further investigate the functional regions of ORF34, a key component of the viral pre-initiation complex (vPIC). These findings are expected to provide more precise insights into the functions and mechanisms of KSHV ORF34 and vPIC. In the realm of life sciences, including virology, deep-learning algorithms have been widely used as powerful tools for predicting protein structure models, and we applied this approach in our early work on a partial vPIC protein-protein interaction model (35). In this study, we derived a structure for KSHV ORF34 using Alphafold2 (AF2), an algorithm capable of predicting protein structure based on amino acid sequences (36). The resulting predicted structure was visualized using an illustration model color-labeled with pLDDT, representing the prediction confidence score (Fig. 1a and 1b). Furthermore, using AF2 and the related module, we visualized the predicted aligned error (PAE) plot, which provides insights into domain prediction by illustrating the predicted distances between alpha-carbon positions for each amino acid pair (Fig. 1c). The green pixels in the PAE plot represent the predicted distance errors between the alpha carbons of the amino acid pairs. Consequently, the presence of green signal areas indicates clustering of amino acid residues that likely form distinct domains. Based on our KSHV ORF34 predicted model, the PAE plot suggested the existence of a small domain in the N-terminal region (approximately 1-50 a.a.) and a larger domain structure in the C-terminal region (approximately 110-327 a.a.) (Fig. 1c). Additionally, this plot indicated that a disordered region (approximately 50-110 a.a.) connects the N-terminal- and C-terminal domains of ORF34 (Fig. 1c). Disordered regions in proteins are typically characterized by the absence of stable tertiary structures, and thus provide flexible linkages between domains with defined structures. In the illustration model with pLDDT (Fig1a and 1b), the predicted disordered region corresponded to the string and helix (labeled orange–yellow), indicating low prediction reliability.

**Fig. 1.**
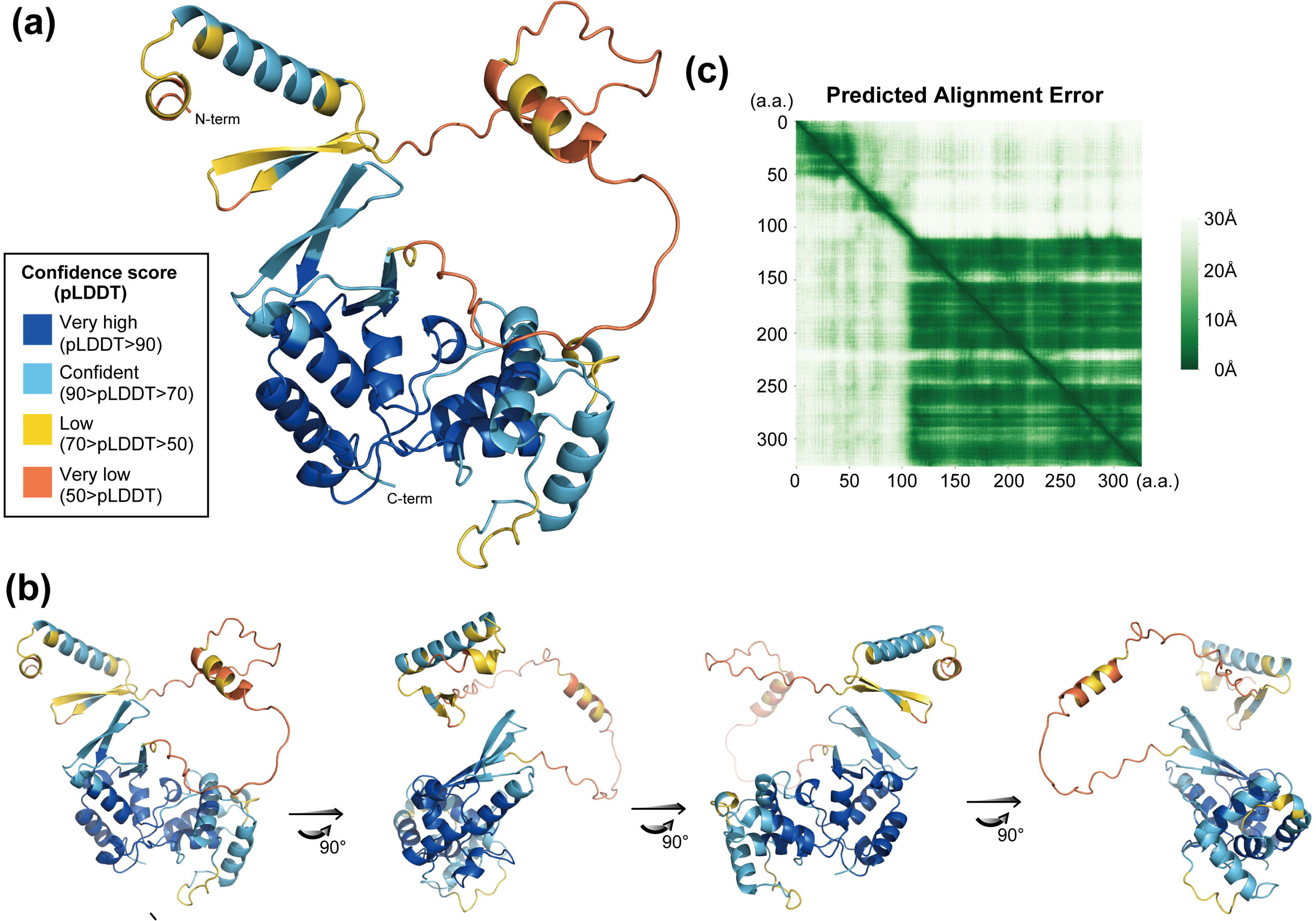
The ORF34 structure model predicted by Alphafold2 and conserved cysteines on ORF34. The predicted ORF34 structure was visualized, and the predicted domain structure was analyzed. (a) The ORF34 structure was predicted using the deep-learning algorithm Alphafold2 (AF2). The illustration depicts the protein labeled with colors according to the pLDDT score of each amino-acid residue. (b) Rotation of the predicted ORF34 structure model that was depicted in (a). (c) Prediction alignment error (PAE) plot of the predicted ORF34 structure. Each white to green pixel indicates a predicted distance of, 30 to 0 Å respectively, between amino-acid residues.

As previously described, the vPIC machinery and its viral components are broadly conserved across beta- and gamma-herpesvirus families (24, 26). It has been hypothesized that critical domains or structures in each functioning protein, such as vPIC components, are conserved among viral homologs. In general, these conserved domains and structures often contain conserved amino acid residues. To ascertain the conserved amino acid residues, we performed sequence alignments of the KSHV ORF34 amino acid residues with other herpes virus homologs (Fig. 2a), which highlighted these conserved residues against a gray background. We discovered 7 amino-acid residues at the N-terminus and 21 at the C-terminus of KSHV ORF34 that were conserved among beta- and gamma-herpesviruses. Thus, to analyze the roles of these conserved residues in ORF34 function, we prepared 18 expression plasmids with single or double alanine substitutions at the conserved amino acid residues to generate ORF34 alanine-scanning mutants, after excluding the conserved prolines to avoid disrupting the local and whole protein backbone (Fig. 2a). Based on the relationships between conserved residues and the predicted structure of ORF34, we generated a histogram displaying the pLDDT score against the amino acid residues of ORF34 with dotting on the conserved residues targeted for the alanine mutations described above (Fig. 2b). The histogram confirmed that the local structural features surrounding these conserved residues were predicted with high confidence, meaning that all conserved residues were localized in a highly structured domain, the N-term region, and the C-term region, as suggested (Fig. 1c). We prepared 18 expression plasmids with single or double alanine substitutions at the conserved amino acid residues to generate ORF34 alanine-scanning mutants, after excluding the conserved prolines to avoid disrupting the local and whole protein backbone (Fig. 2).

**Fig. 2.**
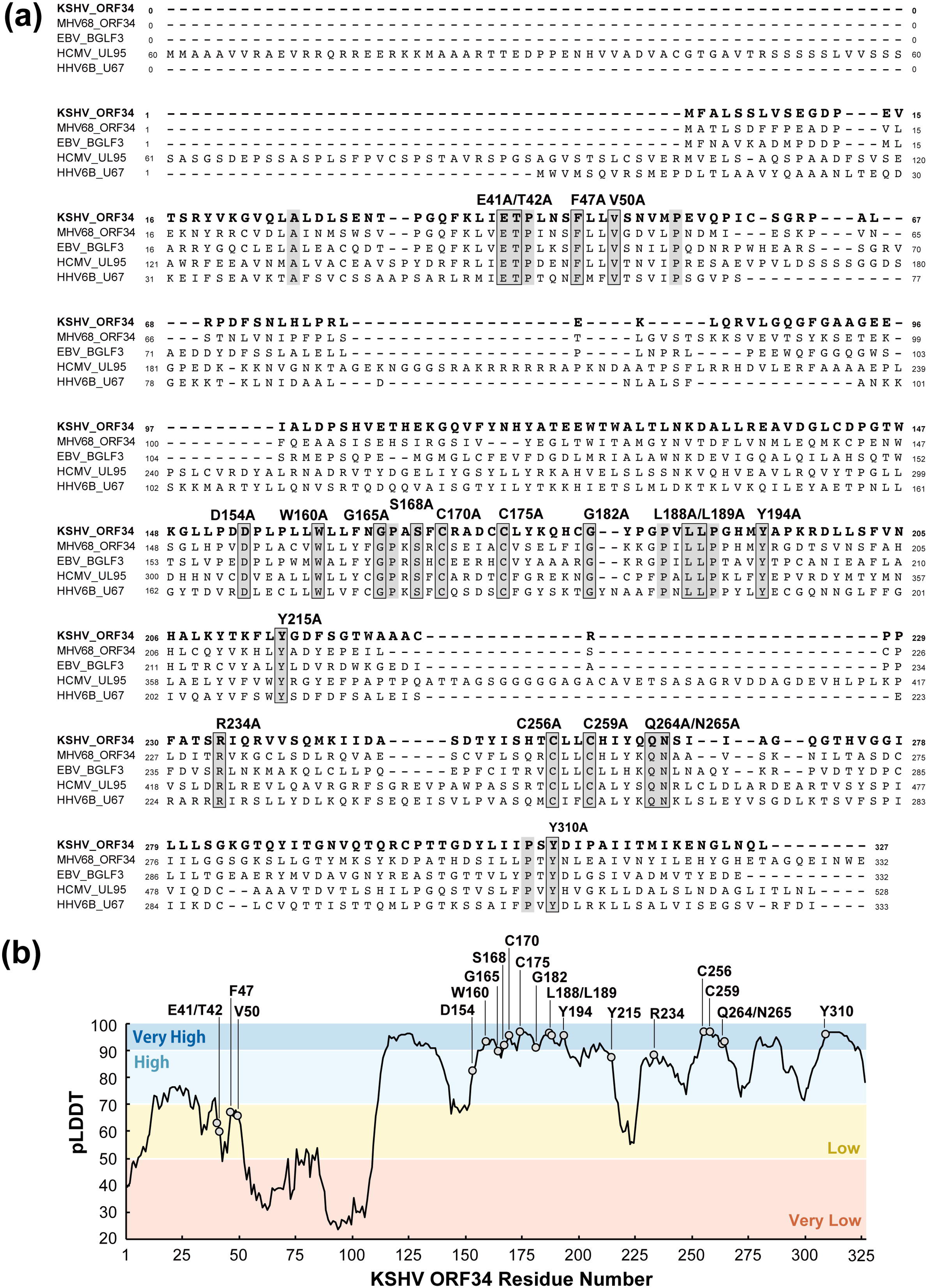
Conserved amino-acid residues of ORF34 homolog and alanine-scanning mutants. Amino acid sequence alignment of KSHV ORF34. Herpesvirus homolog amino acid sequences were translated from nucleotide sequences found in the NCBI database (KSHV ORF34 (JSC-1-BAC16; Accession number GQ994935), MHV68 ORF34 (strain WUMS; NC_001826), EBV BGLF3 (strain B95-8; V01555), CMV UL95 (strain Towne; FJ616285), HHV6B U67 (strain japan-a1; KY239023)). Raw data regarding alignment were obtained using Clustal Omega (EMBL-EBI; https://www.ebi.ac.uk/Tools/msa/clustalo/). Completely conserved amino acids between homologs are indicated by a gray background. Based on the alignment information, 1-2 conserved amino acids residues, except for proline and alanine, were substituted with alanine for ORF34 alanine-scanning mutants, as depicted. (b) The histogram of the pLDDT score demonstrated by AF2 on each amino-acid residue on ORF34 (1-327 a.a.), the conserved amino-acid residues are indicated with white circles.

### Physical interactions between ORF34 alanine-scanning mutants and vPIC components

KSHV ORF34 has been reported to directly interact with the majority of vPIC components (ORF18, ORF24, ORF31, and ORF66), with the exception of ORF30 (27, 28). Based on these findings, KSHV ORF34 is thought to be a hub protein that forms vPIC. To analyze the contribution of ORF34 conserved residues to the interaction with other vPIC components, binding between ORF34 alanine-scanning mutants and the other components was assessed through pull-down experiments (Figs. 3 and 4).

**Fig. 3.**
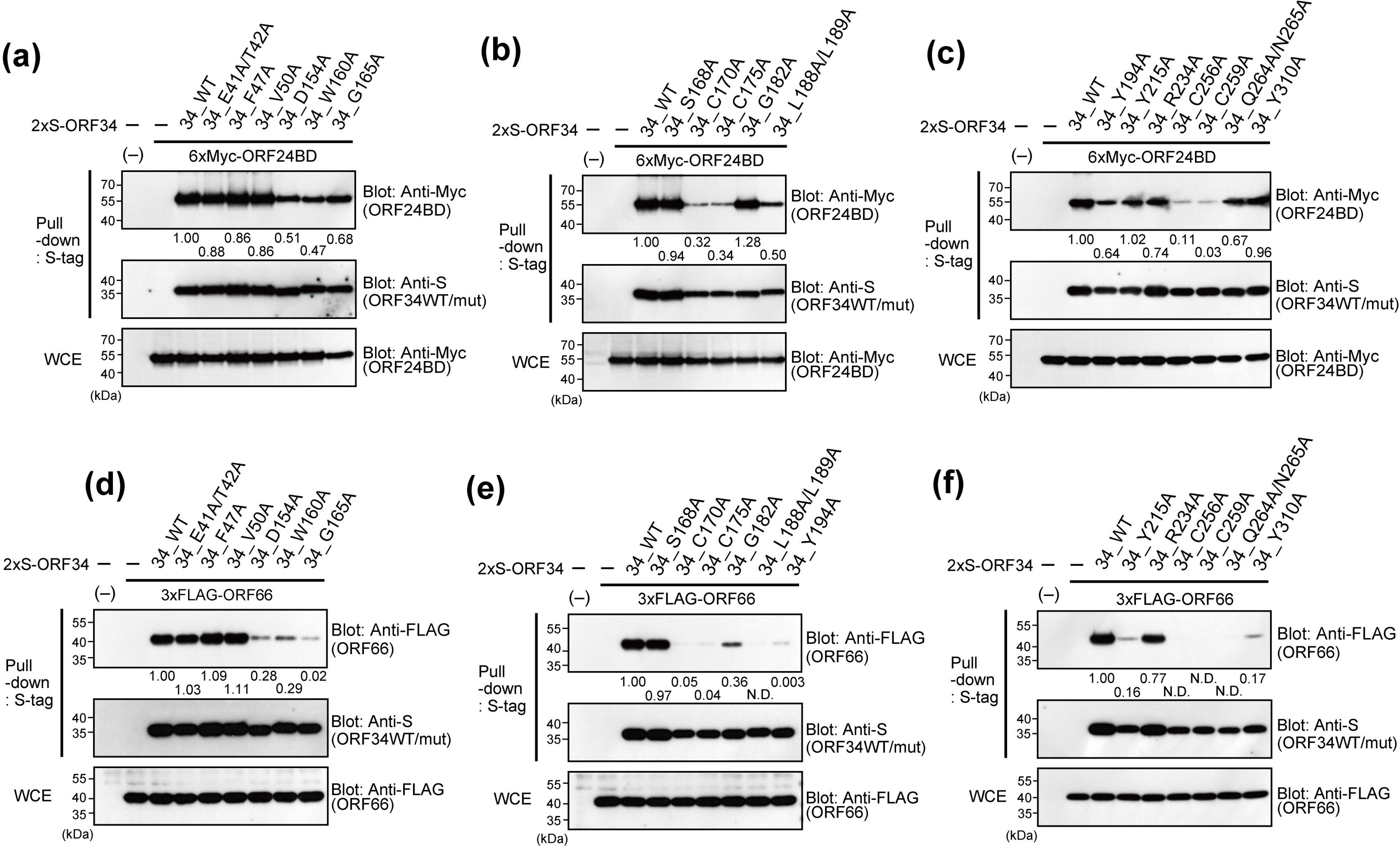
Physical interactions between ORF34 alanine-scanning mutants and vPIC components, ORF24 and ORF66. 293T cells were co-transfected with expression plasmids of the 2xS-ORF34 alanine-scanning mutant and 6xMyc-ORF24BD (a-c) or 3xFLAG-ORF66 (d-f). ORF24BD corresponds to the KSHV ORF24 N-terminal domain (1-400 a.a.) reported as having a domain interaction with ORF34 (28). Transfected cells were lysed, and cell lysates were subjected to pull-down assays using S-protein-agarose that captured the 2xS-ORF34 mutants. Obtained precipitates including 2xS-ORF34 mutants were probed with indicated antibodies to detect interactions. Whole cell extracts (WCEs) were also probed as the input controls of prey proteins for each sample. The pull-down efficiency score of each interaction was indicated below the pull-down panel of prey proteins (6xMyc-ORF24BD or 3xFLAG-ORF66). The pull-down efficiency score of the sample co-transfected with ORF34 WT and ORF24BD/ORF66 was defined as 1.0. N.D. (not detected) indicates that a positive signal of the prey proteins was not detected.

**Fig. 4.**
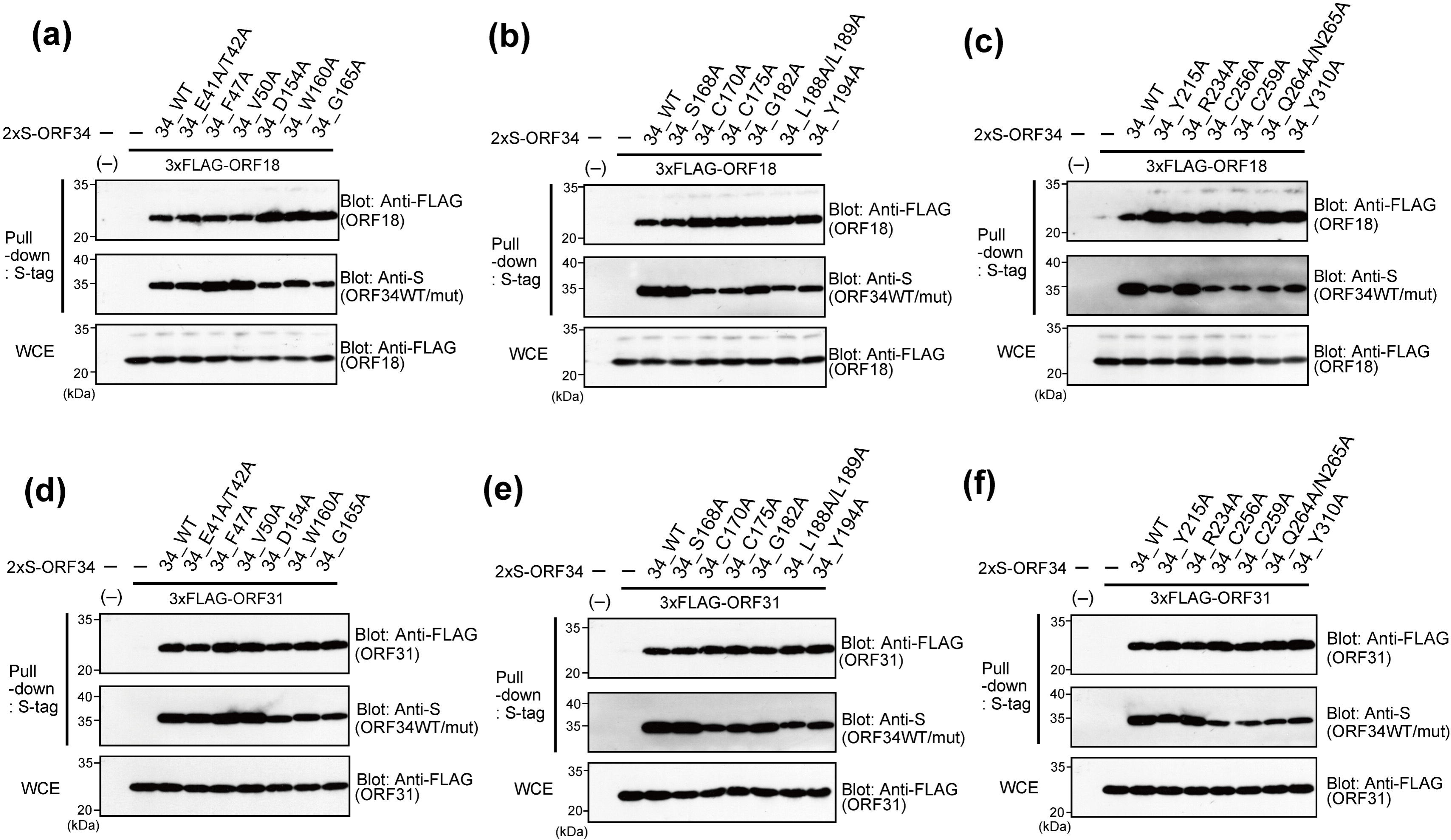
Physical interaction between ORF34 alanine-scanning mutants and vPIC components, ORF18 and ORF31. 293T cells were co-transfected with expression plasmids of 2xS-ORF34 alanine-scanning mutant and 3xFLAG-ORF18 (a-c) or 3xFLAG-ORF31 (d-f). Transfected cells were lysed and cell lysates were subjected to pull-down assays using S-protein-agarose that captured the 2xS-ORF34 mutants. Obtained precipitates including 2xS-ORF34 mutants were probed with the indicated antibodies to detect interactions. Whole cell extracts (WCEs) were also probed as the input controls of prey proteins for each sample.

The N-terminal region (1-400 a.a.) of ORF24 (ORF24 binding domain; ORF24 BD) binds to ORF34, with R328 on ORF24 being essential for ORF34 binding (28). To facilitate ORF24 detection, we prepared expression plasmids of ORF24 BD (6xMyc-tagged ORF24 BD [1-400 a.a.]). 293T cells were co-transfected with 2xS-tagged ORF34 alanine-scanning mutants and 6xMyc-ORF24 BD, 3xFLAG-ORF66, 3xFLAG-ORF18, and 3xFLAG-ORF31, as indicated in each figure, and cell extracts were then subjected to pull-down experiments. Initially, we profiled the interaction ability of the ORF34 mutants with ORF24 BD and ORF66. Our ORF34 mutants exhibited various features associated with another vPIC component. Notably, ORF34 C170A, C175A, C256A, and C259A remarkably decreased or lost their binding abilities to ORF24 and ORF66 (Fig. 3b-c and 3e-f). In addition, G165A, L188A/L189A, Y194A, and Q264A/N265A also lost their interactions with ORF66 (Fig. 3d-f). Conversely, no defects in the interaction between ORF18 and ORF31 were observed in any of the ORF34 alanine-scanning mutants (Fig. 4). Our previous study using ORF34 deletion mutants showed that the 100-150 a.a. region of ORF34 is responsible for binding to ORF18 and ORF31 (27). However, the ORF34 alanine-scanning mutants in the present study did not include any alanine-substituted residues in that region. Therefore, the binding data for the ORF34 alanine-scanning mutants in the present study yielded concordant results.

### Characterization of ORF34 conserved residues concerning KSHV production

Because transcriptional initiation through the vPIC is essential for KSHV production, we considered progeny virus production to be the most appropriate method for evaluating the functional importance of ORF34 conserved residues in vPIC. Therefore, a complementation assay was conducted using ORF34-deficient virus-producing cells stably expressing ORF34 alanine-scanning mutants.

First, we established iSLK cell lines harboring KSHV ΔORF34-BAC (iSLK-Δ34) and iSLK cell lines harboring KSHV ΔORF34 revertant-BAC (iSLK-Δ34Rev). BACmids were constructed as described in our previous study (27). 3xFLAG-tagged ORF34 WT were transfected with the iSLK-Δ34 cells. The control plasmid was transfected into iSLK-Δ34 and iSLK-Δ34Rev cells. To generate stable cell lines, the transfected cells were subjected to drug selection, and complemented exogenous ORF34 in iSLK-Δ34 cells was confirmed (Fig. 5a). Using these cell lines, we evaluated the recovery ability of exogenous ORF34 in iSLK-Δ34 cells (Fig. 5b) and cultured them in conditioned medium containing sodium butyrate (NaB; an HDAC inhibitor that accelerates gene expression) and doxycycline (Dox, an activator of the Tet-on system) for 3 days. Using this system in an iSLK cell, Dox induced ORF50/k-RTA, thereby triggering the lytic phase). The encapsidated KSHV genome was purified from the cultured supernatant and quantified by quantitative polymerase chain reaction (qPCR).

**Fig. 5.**
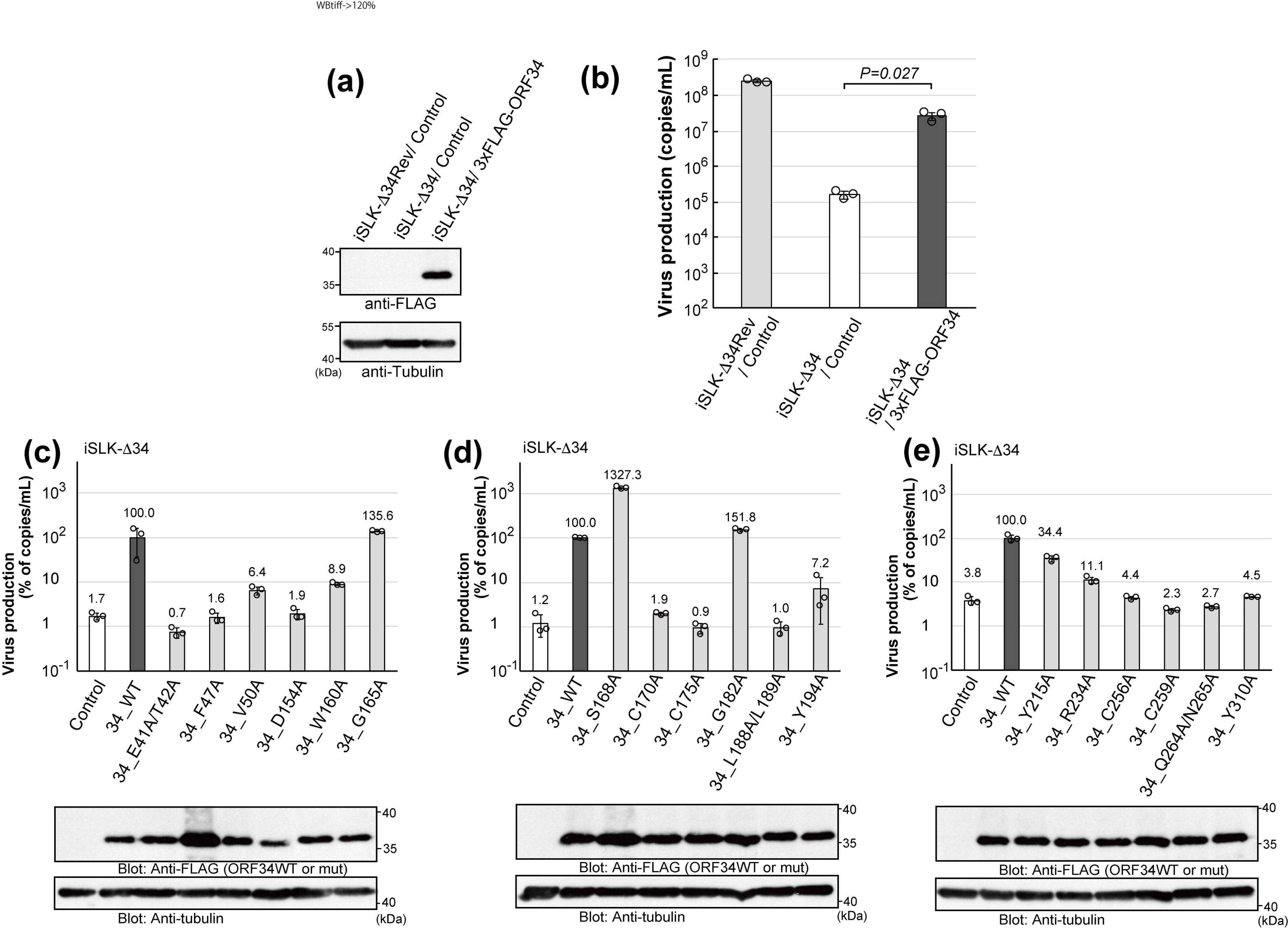
Complement abilities of ORF34 alanine-scanning mutants on viral production. Establishment of 3xFLAG-tagged ORF34-expressing iSLK-Δ34 stable cell lines and confirmation of the recovery by ORF34 WT in ΔORF34-virus production. (a) The Western blot showed exogenous 3xFLAG-tagged ORF34 WT expression in iSLK-Δ34 cells. (b) A complementation assay was performed to rescue virus production. (c-e) 3xFLAG-tagged ORF34 alanine-scanning mutant-expressing iSLK-Δ34 stable cell lines were investigated for their virus production recoveries, with results shown in the upper graph panels. The lower blot panels indicate the exogenous ORF34 WT/mutant expression under normal culture conditions. In the upper graph panels, the virus production of iSLK-Δ34/ORF34 WT cells was defined as 100%. Each stable iSLK-KSHV BAC-harboring cell line was activated by NaB (0.75 mM) and Dox (4 μG/mL) and cultured for 3 days, and the culture supernatant containing encapsidated KSHV genome was harvested and quantified. Three independent samples were evaluated by real-time PCR and indicated as dots in each column. The error bars indicate standard deviations.

The results showed that ORF34 deficiency causes a notable decrease in virus production, and exogenous ORF34 in deficient cells complements most virus production. These results underscore the need for ORF34 in KSHV viral production.

The same strategy was employed to evaluate 18 conserved amino acid mutants of ORF34. Each FLAG-tagged ORF34 mutant expression plasmid was transfected into iSLK-Δ34 cells and selected for the generation of stable cell lines, as indicated, and protein expression was confirmed (lower panels in Fig. 5c-e). KSHV production in these cell lines was then compared with that in iSLK-Δ34/ORF34 WT cells (upper graphs in Fig. 5c-e). The results showed that complementation with ORF34 G165A, S168A, and G182A rescued virus production in iSLK-Δ34 cells, indicating that these substituted residues do not contribute to the ORF34 role in virus release. In contrast, ORF34 E41A/T42A-, F47A-, D154A-, C170A-, C175A-, L188A/L189A-, C256A-, C259A-, Q264A/N265A-, and Y310A-expressing cell lines did not show recovered virus production, indicating that these mutated residues contribute to the ORF34 wild-type role in virus release. Exogenous ORF34 V50A, W160A, Y194A, Y215A, and R234A partially complemented the loss of ORF34 in the iSLK-Δ34 cells.

### Role of ORF34 conserved residues in viral gene transcription and protein expression

The main function of vPIC in the KSHV life cycle is to initiate L-gene expression. To determine whether the virus production recovery abilities of ORF34 WT and mutants were due to KSHV L-gene expression, we analyzed the expression of the representative genes in each step (*i.e.,* K4 as IE, ORF6 as E, and K8.1 as L-gene) based on the mRNA (Fig. 6a-c) and protein expression (Fig. 6d-f). The stable cell lines, iSLK-Δ34 cells complemented with ORF34 WT or alanine-scanning mutants, were stimulated with NaB and Dox for 3 days. Total RNA was extracted from each cell line and subjected to reverse transcription quantitative PCR (RT-qPCR).

**Fig. 6.**
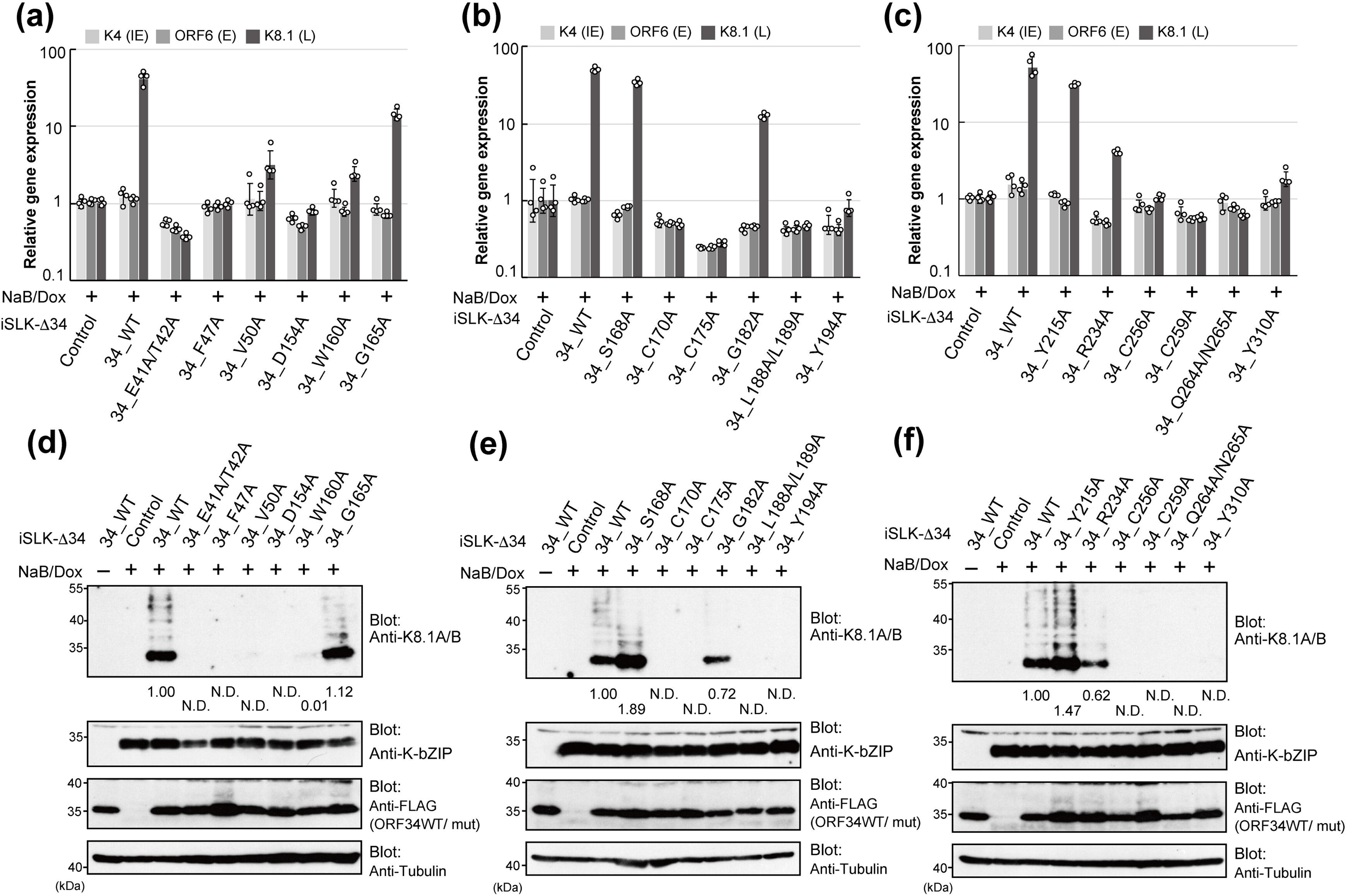
Complement abilities of ORF34 alanine-scanning mutants for viral gene transcription and protein expression. The mRNA and protein expression in lytic-induced iSLK-Δ34 cells complemented with ORF34 WT/alanine-scanning mutants. The cells were cultured for 3 days in conditioned medium containing NaB and Dox to induce a lytic state. (a-c) Total RNA was extracted from cells and subjected to RT-qPCR. The mRNA expression of viral genes, which is representative of each step (K4 as IE gene, ORF6 as E gene, K8.1 as L-gene), was normalized by GAPDH expression. The mRNA expression of iSLK-ΔORF34/control cells with NaB and Dox treatment was defined as 1.0. Four independent samples were evaluated and indicated as dots in each column. The error bars indicate standard deviations. (d-e) Total protein samples were prepared from each lytic-induced cell and subjected to Western blotting. The protein expression of the L-gene was analyzed by K8.1A/B blotting and the E gene by K-bZIP, respectively. The protein expression score, the band intensity, of iSLK-Δ34/ORF34 WT was defined as 1.0. N.D. (not detected), indicating that a positive signal of viral proteins was not detected.

The results showed that the transcription of K8.1 in iSLK-Δ34 cells expressing ORF34 E41A/T42A, F47A, D154A, C170A, C175A, L188A/L189A, Y194A, C256A, C259A, Q264A/N265A, and Y310A was markedly reduced compared with that in iSLK-Δ34/WT cells, but differed little from that in iSLK-Δ34/control cells. In contrast, the transcription of K8.1, complemented by G165A, S168A, G182A, and Y215A (Fig. 6a-c). The transcription levels of K4 and ORF6 were almost equal in all iSLK-Δ34 cell lines expressing ORF34 WT or its mutants. In parallel with RNA sampling, we prepared protein samples from each complemented iSLK-Δ34 cell line. The protein expression of the representative E gene K-bZIP in each cell line showed no marked differences among cell lines (Fig. 6d-f; upper middle panels). In contrast, we observed notable differences in K8.1 protein expression among the cell lines. The K8.1 protein expression was rescued in ORF34 G165A-, S168A-, and Y215A-expressing iSLK-Δ34 cell lines and was partially complemented in ORF34 R234A- and G182A-expressing cells. The protein expression of K8.1 was unable not be rescued in other mutants (Fig. 6d-f; upper panels). These differences in K8.1 protein expression among cell lines were consistent with the data on K8.1 mRNA obtained by qPCR.

### Contribution of ORF34 conserved residues for recruitment to K8.1 transcription start site

vPIC formation at the TSS of L-genes is essential for the expression of the L-genes. To assess the influence of substitution of conserved residues for ORF34 recruitment to K8.1-TSS, we evaluated the association of ORF34 WT and mutants with K8.1-TSS in the lytic state of the KSHV genome. The stable cell lines, iSLK-Δ34 cells complemented with ORF34 WT or alanine-scanning mutants, were stimulated with NaB and Dox for 3 days. The lytic-induced cell lines were subjected to ChIP with Contorol IgG or anti-FLAG antibody and then subjected to qPCR (ChIP-qPCR) (Fig. 7a-c).

**Fig. 7.**
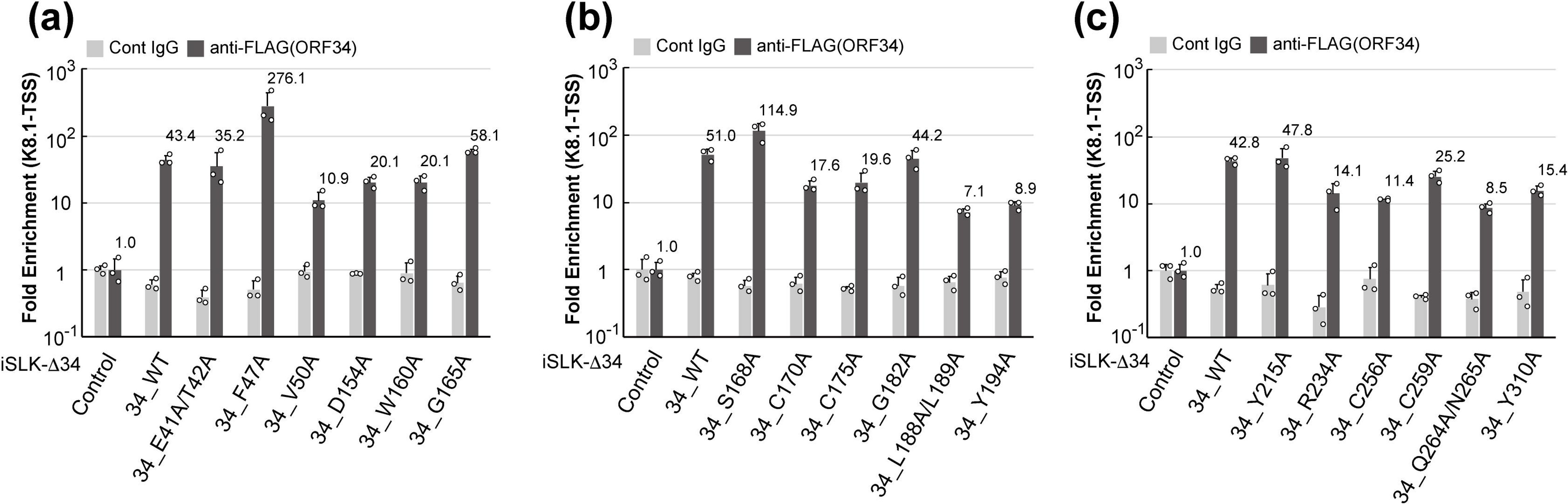
Recruitment of ORF34 alanine-scanning mutants to late gene transcription start-site. The binding quantity of the KSHV late gene TSS to ORF34 WT/alanine-scanning mutants was evaluated by ChIP. The iSLK-Δ34 cells complemented with ORF34 WT/alanine-scanning mutants were cultured for 3 days in conditioned medium containing NaB and Dox to induce a lytic state. (a-c) A KSHV DNA fragment precipitated by control mouse IgG or anti-FLAG monoclonal antibody was subjected to RT-qPCR and normalized by input DNA subjected to ChIP. Each fold enrichment of iSLK-ΔORF34/control cells by control IgG or anti-FLAG antibodies was defined as 1.0. Three independent samples were evaluated and indicated as dots in each column. The error bars indicate standard deviation.

The results demonstrated that the association of ORF34 with K8.1-TSS in iSLK-Δ34 cells expressing ORF34 F47A and S168A was clearly higher than that in iSLK-Δ34/WT cells. The complement abilities of E41A/T42A, G165A, G182A, and Y215A were almost the same as those of the ORF34 WT. The remaining cell lines complemented with the mutants V50A, D154A, W160A, C170A, C175A, L188A/L189A, Y194A, R234A, C256A, C259A, Q264A/N265A, and Y310A, showed a decline in the enrichment of K8.1-TSS in comparison to iSLK-Δ34/ WT cells, but did not reach the basal level (*e.g.*, iSLK-Δ34/control cells) (Fig. 7a-c).

### Profiling of ORF34 alanine-scanning mutants on vPIC-related parameters

The data obtained in this study (Fig. 3, 5-7) were summarized and visualized using a heatmap with a dendrogram of hierarchical clustering (Fig. 8a). The extracted parameters were as follows: i) binding to ORF66 (Fig. 3d-f), ii) binding to ORF24BD (Fig. 3a-e), iii) K8.1 mRNA expression (Fig. 6a-e), iv) K8.1A/B protein expression (Fig. 6d-f), v) KSHV production (Fig. 5c-e), and vi) ORF34 recruitment to K8.1-TSS (Fig. 7a-c). Binding to ORF18 and ORF31 (Fig. 4) was excluded because no differences between ORF34 WT and its mutant were observed. ORF34 mutants were roughly classified into 4 groups according to the visualized dendrogram (Fig. 8a). In parallel, we also calculated the relative solvent-accessible surface area (relative-SASA), the indicator of amino-acid residues exposed on the protein surface, based on the ORF34 prediction model (Fig. 8b). The classified residues are highlighted in the illustration (Fig. 9a) or illustration-surface model of the predicted ORF34 (Fig. 9b).

**Fig. 8.**
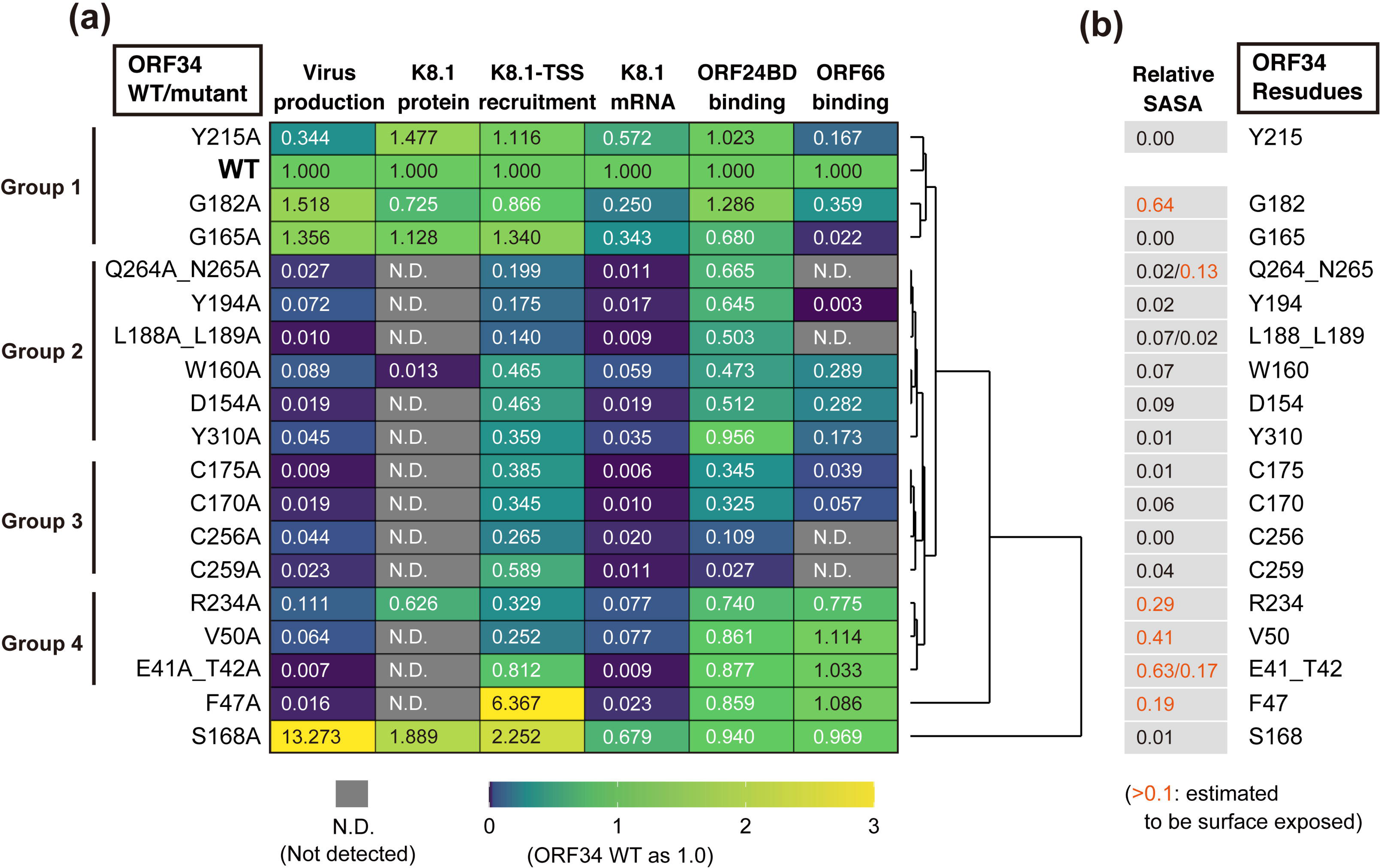
Summary of the effects of ORF34 alanine-scanning mutants on the vPIC function. (a) The results (Fig. 5, 5-7) were summarized and visualized on the heatmap with a dendrogram highlighting the hierarchical clustering. Each score of ORF34 WT or iSLK-Δ34/WT was defined as 1.0. The scores are indicated on each panel with the color gradient (dark blue [0.0] - green [1.0] - yellow [>3.0]) as indicated by the lower bar. N.D. (not detected) in each experiment is indicated by the gray panel. Groups were classified according to the visualized dendrogram. (b) Based on the ORF34 structure model (Fig.1a), the prediction score of protein surface exposure, relative solvent accessible surface area (relative-SASA), were listed for each mutated site. The relative-SASA score colored with orange [>0.1] indicates that the amino-acid residue has a high possibility of being exposed on the protein surface.

**Fig. 9.**
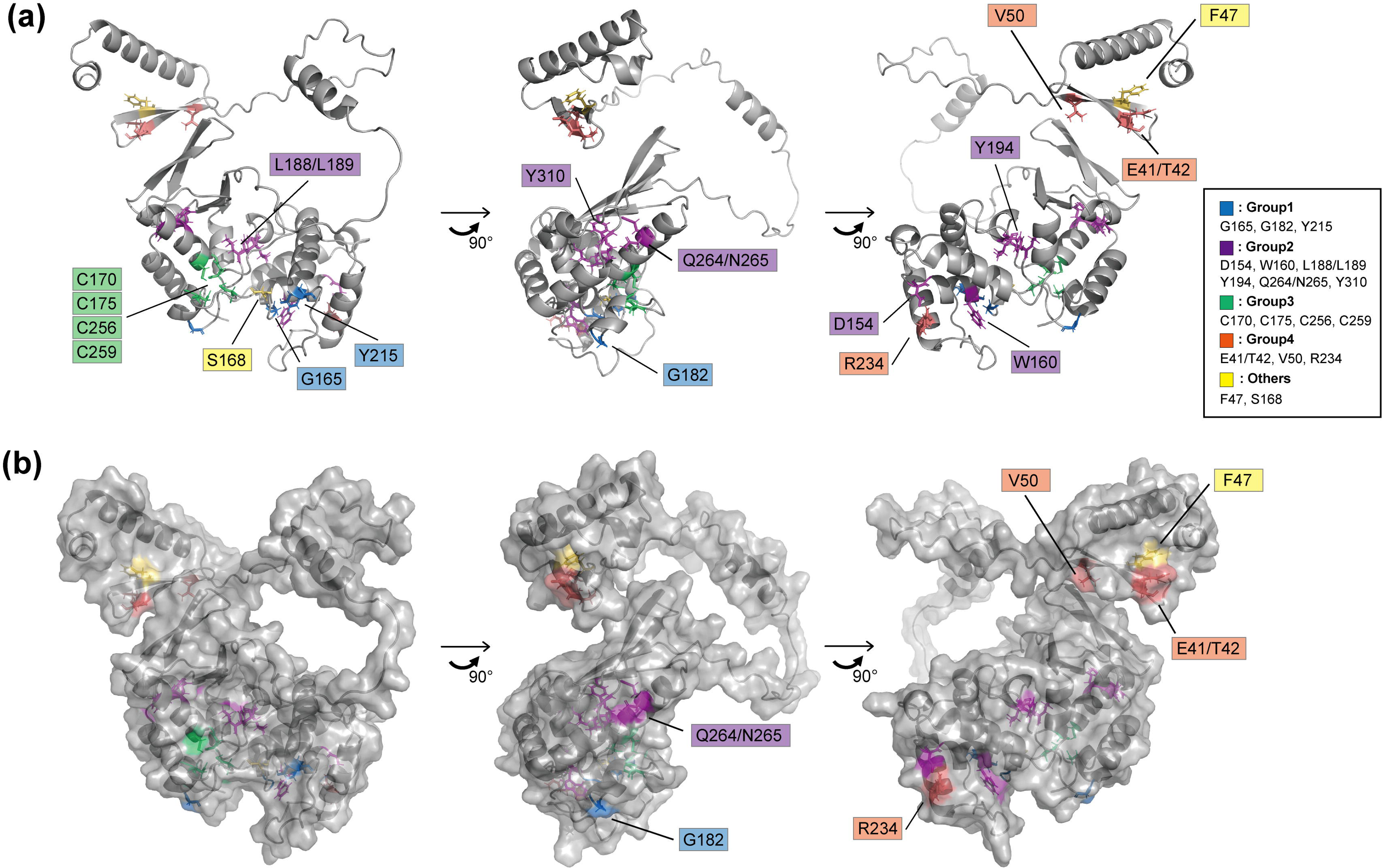
Overview of the predicted ORF34 structure and mutated residues. Rotated ORF34 structures with caption based on the predicted structure (Fig 1a and 1b) (a) In the illustration model, all the mutated amino-acid residues are highlighted with colors according to the grouping (Fig. 8a), and captioned. (b) In the surface-illustration model based on (a), only the mutated amino-acid residues expected to be exposed on protein surface (Fig. 8b) are captioned.

Group 1: Mutants complemented virus production in iSLK-Δ34 cells. Thus, these substituted amino acid residues are not essential for vPIC function.

Group 2: The mutants maintained their binding to ORF24 (but not to ORF66) and failed to complement virus production. These mutated sites may be related to vPIC function via ORF66 binding. It is possible that these mutated sites are related to the vPIC function via ORF66 binding.

Group 3: The mutants failed to complement virus production and showed decreased binding to both ORF24 and ORF66. Conserved residues in the mutants were inferred to be essential for vPIC function via complex formation.

Group 4: While the mutants did not rescue virus production, their binding ability to ORF24 and ORF66 was unaltered. Thus, these mutation sites are important for the vPIC machinery. The mechanism underlying the contribution of these residues to vPIC function might not involve a direct interaction with ORF24 and ORF66. However, the precise details of this process remain unknown.

The S168A and F47A mutants were out of the range of the other mutants in the dendrogram. The ORF34 S168A mutant showed over-complementation in virus production and K8.1-TSS recruitment, approximately 13 and 2.2 times more than ORF34 WT, respectively. Almost equal to ORF34 WT, ORF34 S168A showed binding abilities to ORF24 and ORF66. Therefore, further studies on S168 are necessary to clarify the mechanisms of vPIC regulation. The ORF F47A mutant also demonstrated over-complementation in K8.1-TSS recruitment, namely it was more than 6 times higher than that of ORF34WT. However, the F47A mutant failed to complement L-gene transcription, protein expression, or virus production. The ORF34 F47 residue, present in the predicted N-term domain, may have roles in vPIC-mediated transcription or vPIC assembly, other than the direct association with other vPIC components.

G165A, G182A, and Y215A, classified as Group 1, reduced the interaction with ORF66 and maintained or mildly reduced viral production. Therefore, these sites likely contribute to the association between the2 viral proteins, ORF34 and ORF66. However, considering the small influence on virus production, L-gene expression, and L gene TSS assembly in each KSHV-producing cell line, other factors brought about by viral infection, including vPIC components, might help restore the association between ORF34 G165A/G182A/Y215A and ORF66 in virus-producing cells.

The mutants classified into Group 2, D154A, W160A, L188A/L199A, Y194A, Q264A/Q265A, and Y310A, showed little recovery of virus production, an almost deficient vPIC function, a mild decrease in association with ORF24, and a decrease in ORF66 (Fig. 3-8a). Prediction models suggest that these mutated sites are included in the inner side of the C-terminal domain, and only N265 was exposed to the protein surface (Fig. 8b and Fig. 10). Thus, these mutated residues are responsible for supporting the backbone of the C-terminal domain structure.

**Fig. 10.**
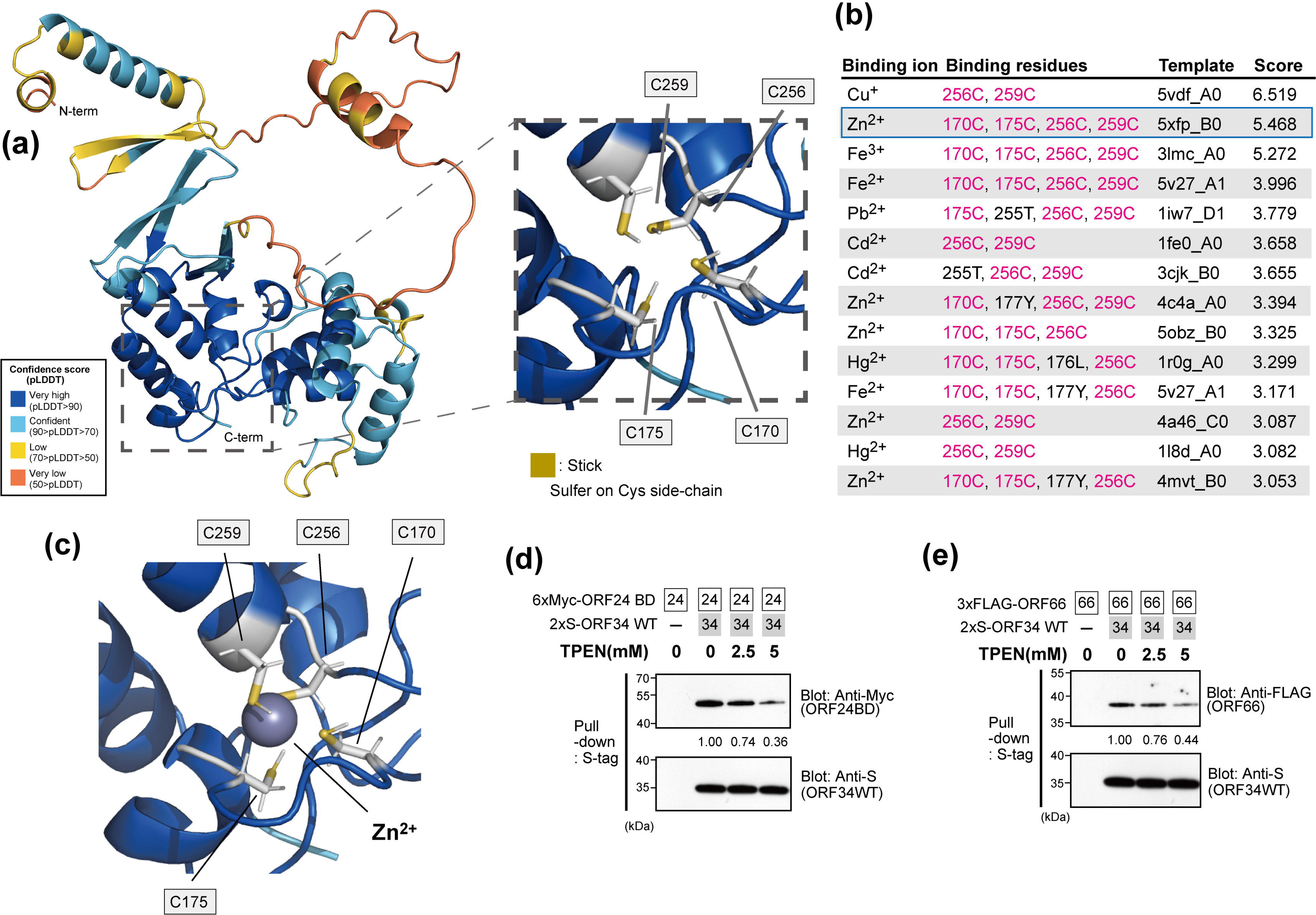
The prediction of metal cation binding to ORF34 and its influence on the binding ability of ORF34. (a) The table organizes the binding residues, templates, and binding scores of 18 different metal ions predicted by the Metal Ion-Binding site prediction and modeling server, MIB2. When multiple copies of identical binding residues were present for a single metal ion, the template with the highest binding score was noted. If any new binding residues were present amongst previously recorded ones, the template was still recorded. Prediction binding scores higher than 3.0 are listed. Conserved cysteine residues (C170, C175, C256, C256) were highlighted with magenta. (b) Predicted model of ORF34 structure binding to ions. The illustration model of ORF34 was visualized according to Fig. 7a (right panel), and the binding zinc ion (Zn^2+^) is shown as a gray sphere. Heavy metal cation chelation influences the ORF34 binding abilities to ORF24BD (c) and ORF66 (d). Pull-down assays using S-tagged ORF34 binding beads as bait in the presence of cell extracts overexpressed prey proteins (6xMyc-ORF24BD or 3xFLAG-ORF66) and the heavy metal chelator TPEN. The pull-down efficiency score is indicated below the pull-down panel of the prey protein. The score of the sample co-incubated with 2xS-ORF34 WT binding beads as bait and 6xMyc-ORF24BD or 3xFLAG-ORF66 as prey was defined as 1.0.

All ORF34 mutants, which maintained a minimum binding ability to ORF24, were enriched in the L-gene TSS to some extent (Fig. 7a-c and 8a), although their expression was not fully correlated with the degree of association with ORF24BD (Fig. 3a-c and 8a). However, the mutants in groups 2, 3, and 4 failed to recover viral production. This suggests that the mutated residues of groups 2 and 3 contribute to the vPIC function for virus production via an association with ORF66, and mutated residues of group 4 operate in vPIC via other mechanisms.

At a minimum, our data (Fig. 3, 5-7) and grouping of ORF34 mutants by hierarchical clustering (Fig. 8a) demonstrated that ORF34 C170A, C175A, C256A, and C259A, which were categorized into Group 3, lost most of their interactions with ORF24 and ORF66 and the virus production recovery ability. This suggests that these conserved cysteines, which were predicted to have little exposure to the protein surface (Fig. 8b and 9a-b), are crucial for ORF34 function in KSHV replication.

### Four conserved cysteine residues in the predicted ORF34 structure

Among the conserved amino acids, 4 cysteines (C170, C175, C256, and C259) were predicted to be present in the C-terminal domain (Fig. 1c and 2b), including the inner side of the protein (Fig. 8b and 9). The precise position and side chain were visualized and enlarged (Fig. 10a). On reframing these residues in an amino acid sequence (Fig. 2a), C170-R171-A172-D173-C174-C175 and C256-L257-L258-C259 form the consensus motif sequences C-X_4_-C and C-X_2_-C, respectively. These 2 consensus sequences (C-X_n_-C) are known to form ion-capturing domains such as the zinc finger domain. Furthermore, in the prediction model, we found that the pairs of consensus C-X_n_-C motifs exhibited nearly symmetrical positioning within ORF34 (Fig. 10a), suggesting the potential for metal cation capture.

To simulate the ion-binding affinity resulting from the conserved cysteine residues in our ORF34 structural model, we estimated potential binding ions by employing a Metal Ion-Binding site prediction and modeling server (37). Docking simulation predicted the binding residues of ORF34 with 16 metals and 18 ions. We identified metal ions that were captured by residues within at least one of the consensus sequence cysteines (C170-C175 and/or C256-C259) and had predicting scores > 3.0 (Fig. 10b). Of the 18 ions tested, 7 (Zn^2+^, Fe^2+^, Fe^3+^, Cd^2+^, Hg^2+^, Cu^+^, and Pb^2+^) were predicted to be captured by at least 1 of the C-Xn-C ORF34 motifs. Notably, 3 metal ions (Zn^2+^, Fe^2+^, and Fe^3+^) were expected to be captured by all 4 conserved cysteine residues (Fig. 10b), with the binding score indicating a dominant interaction between ORF34 and Zn^2+^ ions.

Next, we attempted to confirm that metal ion capture is crucial for ORF34 to function as a hub protein in vPIC. The *in vitro* binding ability of ORF34 to ORF24 or ORF66 was analyzed in the presence of the chelating agent, TPEN, which has a higher affinity for heavy metal ions, including Zn, Fe, and Mn, than for Ca and Mg. The stability constant, a chemical indicator of chelating ability, of TPEN was the highest for Zn among the 5 metal ions. Our results showed that TPEN interferes with ORF34 association not only with ORF24 BD (Fig. 10c) but also with ORF66 (Fig. 10d) in a concentration-dependent manner. These results suggested that capturing the heavy metal cation via conserved cysteines contained in the C-X_n_-C consensus motif is important for the binding of ORF34 to other vPIC components (ORF24 and ORF66).

## Discussion

In this study, we constructed alanine-substituted mutants of KSHV ORF34 with conserved amino acid residues and assessed their roles in viral replication and vPIC formation. After obtaining diverse information from our results, it is noteworthy that the 4 conserved cysteines (C170, C175, C256, and C259) in KSHV ORF34 are essential for the vPIC machinery. The mutants KSHV ORF34 C170A, C175A, C256A, and C259A were deficient in their association with ORF24 and ORF66 (Fig. 3 and 8a). Furthermore, these mutants failed to complement virus production (Fig. 5 and 8a) and K8.1 gene transcription and protein expression (Fig. 6 and 8a) but also reduced recruitment to K8.1-TSS (Fig.7 and 8a). The conserved cysteines of ORF34 may have some physical or virological significance in protein structure and function. One of the findings supporting this speculation is that the 4 cysteines comprised 2 pairs of C-X_n_-C consensus sequences (C170 and C175, and C256 and C259) (Fig. 2a). Second, the deep-learning-structure model predicted that the conserved cysteines are present on the inner side of ORF34 (Fig. 8b, 9a-b, and 10a) and are placed in a roughly tetrahedral formation (Fig. 9 and 10a). Using the AlphaFold2 prediction model as our query, another prediction method (Metal Ion-Binding site prediction and modeling server; MIB2) provided data supporting the high probability of ion holding by ORF34 conserved cysteines (Fig. 9b). This prediction demonstrated that the top candidate metal cation captured by the double pair of conserved cysteines of ORF34 was zinc (Fig. 9b).

Further supporting the speculation concerning the physical or virological significance of the conserved cysteines, the binding ability of ORF34 to ORF24 and ORF66 was decreased in the presence of TPEN, a heavy metal cation chelator (Fig. 9c and 9d). Because TPEN has a higher affinity for zinc cations than iron, it can exclude captured zinc more efficiently than iron and other metal cations. An article regarding the mathematical prediction of the ion-binding site of metalloproteins revealed that AlphaFold2 is a useful and extensive tool for protein annotation of zinc ion binding (38). This report endorses our analysis strategy for annotating the 4 conserved cysteines of ORF34 as binding sites for metal cations. Despite the lack of direct evidence, our prediction models and data from chelator treatment suggest that zinc was captured by ORF34 conserved cysteines. Concrete evidence concerning protein structure, including the ion positions as determined by X-ray crystallography, nuclear magnetic resonance (NMR), or Cryogenic Electron Microscopy (Cryo-EM), is indispensable for confirming these findings.

For host proteins composing cellular PIC, the functional analog of vPIC, capturing the ions is also important. Cellular PIC consist of TBP, GTFs, and multiple subunits of the RNAPII complex. Holding zinc via C-X_n_-C motifs has been reported in several factors of cellular PIC, including TFIIEα (39), TFIIB (40), TFIIS (41), and RPB9 (RNAPII subunit 9) (42). In TFIIEα, the loss of capturing ions induces a structural disturbance that negatively influences its binding ability to other factors and its transcription capability (39, 43). Zinc binding to TFIIB affects the structural rigidity and protein surface charge around the zinc-binding domain (44). Furthermore, the C-X_n_-C motif is indispensable for other vPIC components. We and Didychuck *et al.* demonstrated the importance of the C-X_n_-C sequence of ORF66 in viral replication via vPIC formation (30, 31). In summary, the present findings and previous reports of host cellular PIC factors strongly support the notion that ion capture via the C-X_n_-C motif on the KSHV ORF34 protein is crucial for maintaining the structural character and binding ability of ORF34 to other vPIC components, resulting in virus production via L-gene expression using vPIC machinery.

Our functional profiling of the ORF34 conserved residues provided other valuable findings. KSHV ORF34 conserved residues E41/T42 and V50, which were classified into Group 4 (Fig. 8a), and F47, which were not classified (Fig. 8a), were present in the ORF34 N-terminus (fig.2a). E41/T42 and V50 were essential residues for viral replication via L-gene expression but did not affect associations with other vPIC components (Fig. 3-8a). F47 had almost the same tendency as other N-terminal residues, but F47A showed a significant increase in the K8.1 promoter. A simple speculation concerning the roles of these residues is that they are associated with unknown host factors, which contribute to the modulation of vPIC-mediated transcription. The predicted algorithm AlphaFold2 suggested that ORF34 has an N-terminal domain and a C-terminal domain separated by the disordered region (Fig. 1a-b and 2b). E41/T42, F47, and V50 exist in the N-terminal domain (Fig. 2b and 9a-b). The other conserved amino acid residues, including those responsible for binding to other vPIC components, ORF24 and ORF66, exist in the C-terminal domain (Fig. 2b, 8a, 9a-b). Other prediction algorithms for the disordered region, PrDOS (45) and DISOPRED3 (46), also suggested the presence of disordered regions of KSHV ORF34 at 64-70/89-108 a.a. and 92-107 a.a., respectively. Homology search results using the PDB database in the running process of AlphaFold2 demonstrated a similarity between the N-terminal domain and some regulatory factors of the RNAPII complex.

Another hypothesis concerning the conserved N-terminal residues is based on EBV studies. It was reported that the phosphorylated T42 residue of BGLF3 (KSHV ORF34 homolog) is essential for vPIC function and that phospho-T42 of BGLF3 is important for sub-complex formation among BFRF2 (KSHV ORF66 homolog) and BVLF1 (KSHV ORF18 homolog) (47). These reports demonstrated that 2 molecular interactions, BGLF3 binding to BFRF2 or BVLF1, were not affected by the phosphorylation state of T42, but 3 molecular interactions, the presence of BGLF3, BFRF2, and BVLF1, were positively dependent on phospho-T42 (47). Our alanine mutant, ORF34 E41A/T42A, harbors a mutation in analogous residues of EBV BGLF3 T42. Our observations of the ORF34 E41A/T42A mutant, including interaction analysis, gene expression, and virus production, were well matched with a previous EBV study (47). Unfortunately, whether the subcomplex (*i.e.,* a subset of a larger functional protein complex) is present in KSHV has not yet been confirmed. Phospho-T42 in KSHV ORF34 may contribute to subcomplex formation, and the residues surrounding T42 (E41, F47, and V50) may also work in coordination. However, further research on this topic is required.

Another characteristic finding was the conservation of serine and tyrosine residues in the KSHV ORF34. First, the ORF34 serine residue mutant S168A showed a better ability for virus production than ORF34 WT (Fig.5 and 8a). Previous studies on vPIC factors among herpesviruses have identified multiple serine/threonine residues in EBV BDLF4 (KSHV ORF31 homolog) (48). If phosphorylation occurs on KSHV ORF34, S168 phosphorylation is related to the dysregulation of vPIC. Second, the ORF34 tyrosine residue mutants, Y194A and Y310A, lost their ability to bind to ORF66, L-gene expression, translocate to the L-gene TSS, and virus release, but did not lose their ability to bind to ORF18, ORF24, and ORF31 (Fig. 3-8a). Thus, phosphorylation of Y194 and Y310 may be related to the positive regulation of ORF66 binding. However, the prediction model did not indicate the full exposure of the side chains of these residues, S168, Y194, and Y310, to the surface of ORF34 (Fig. 8b and 9a-b). To determine whether phosphorylation of ORF34 occurs, real experimental data are necessary. Phosphorylation and other post-translational modifications might also contribute to vPIC regulation.

Our results demonstrate the importance of the 4 conserved cysteine residues in the KSHV ORF34 protein for functional vPIC, following KSHV ORF66 (30, 31). Our evidence and structure prediction suggest that these 4 cysteines would contribute to cation capturing, which might be essential for maintaining the conformation of ORF34 as a scaffold for vPIC formation. In addition, we can speculate about the functional possibilities of the character profiling of the ORF34 mutants and the predicted structure. The N-terminal domain of ORF34 may be associated with unknown host factors and may bundle multiple vPIC components. On the other hand, according to our prediction model, our experimental strategy using conserved residue mutations would not fully cover the associated protein surface. Thus, adequate proof has not yet been obtained, and further efforts are necessary to answer questions concerning this complex.

## Material and Methods

### Protein structure prediction and ion binding prediction and modeling

The KSHV ORF34 protein structure model was predicted using AlphaFold 2.2.0 software program (https://github.com/deepmind/alphafold) in the local environment (36). The following databases were used for AlphaFold2: Uniclust30 (version 2018_08), MGnify (version 2018_12), pdb70 (downloaded on Feb 15th, 2022), PDB/mmCIF (downloaded on Feb 18th, 2022), and pdb_seqres (downloaded on Feb 18th, 2022). Notably, several versions or frequently updated versions of these databases have been utilized. The predicted model was visualized using the molecular visualization open-source software program PyMOL (ver. 2.5.0). In addition, the pLDDT score coloring of the predicted model was visualized with the python module “PSICO” (https://github.com/speleo3/pymol-psico). Visualization of the PAE plot was completed using the Python module “plddt2csv” (https://github.com/CYP152N1/plddt2csv). To predict disordered protein regions of KSHV ORF34, PrDOS (https://prdos.hgc.jp/cgi-bin/top.cgi) and DISOPRED3 (http://bioinf.cs.ucl.ac.uk/psipred) have also been used (45, 46). The relative per-residue solvent-accessible surface area was calculated using the ‘get_sasa_relative‘ syntax in open-source PyMOL 2.1, with default settings of dot density of 2 and solvent radius of 1.4.

The metal ion-binding (MIB) site prediction and modeling server (http://bioinfo.cmu.edu.tw/MIB/) was used to predict the binding residues of 18 different metal ions (Ca^2+^, Cu^+^, Cu^2+^, Fe^2+^, Fe^3+^, Mg^2+^, Mn^2+^, Zn^2+^, Cd^2+^, Ni^2+^, Hg^2+^, Co^2+^, Au^+^, Ba^2+^, Pb^2+^, Pt^2+^, Sm^3+^, and Sr^2+^) in the protein (37, 49). The query of the protein structure was compared to metal-ion binding templates, which allowed MIB residues within at least 3.5 Å of the ion to be predicted, along with a binding score and template. Template with the highest binding score (> 3.0) was documented if the same binding residue was present multiple times for a single metal ion.

### Cell culture and reagents

293T cells (RCB2202; RIKEN BioResource Center, Tsukuba, Japan) were cultured in the growth medium. DMEM was supplemented with 10% fetal calf serum and a penicillin-streptomycin solution (Nacalai Tesque Inc., Kyoto, Japan). The transfection reagent for 293T cells, PEI-MAX MW40000 (Polysciences, Inc., Warrington, PA, U.S.A.), was dissolved at a concentration of 1 mg/mL in distilled water and filtered.

### Plasmids

The pCI-neo-2xS-ORF34 WT, pCI-neo-3xFLAG-ORF66, pCI-neo-3xFLAG-ORF18, and pCI-neo-3xFLAG-ORF31 expression plasmids have been described previously (27). Alanine scanning mutants of ORF34 coding fragments were obtained by PCR or overlap-extension PCR from the KSHV ORF34 expression plasmid using the primer sets indicated in Table 1. To construct pCI-neo-6xMyc-ORF24-BD (1–400), partial ORF24 coding fragments (1-400 a.a.) containing the ORF34 binding region (28) were also obtained by PCR from the KSHV ORF24 expression plasmid. These fragments were cloned into expression plasmids based on a pCI-neo mammalian expression vector (Promega, Madison, WI, U.S.A.), *e.g.* pCI-neo-2xS, pCI-neo-6xMyc, or pCI-blast-3xFLAG.

**Table 1:**
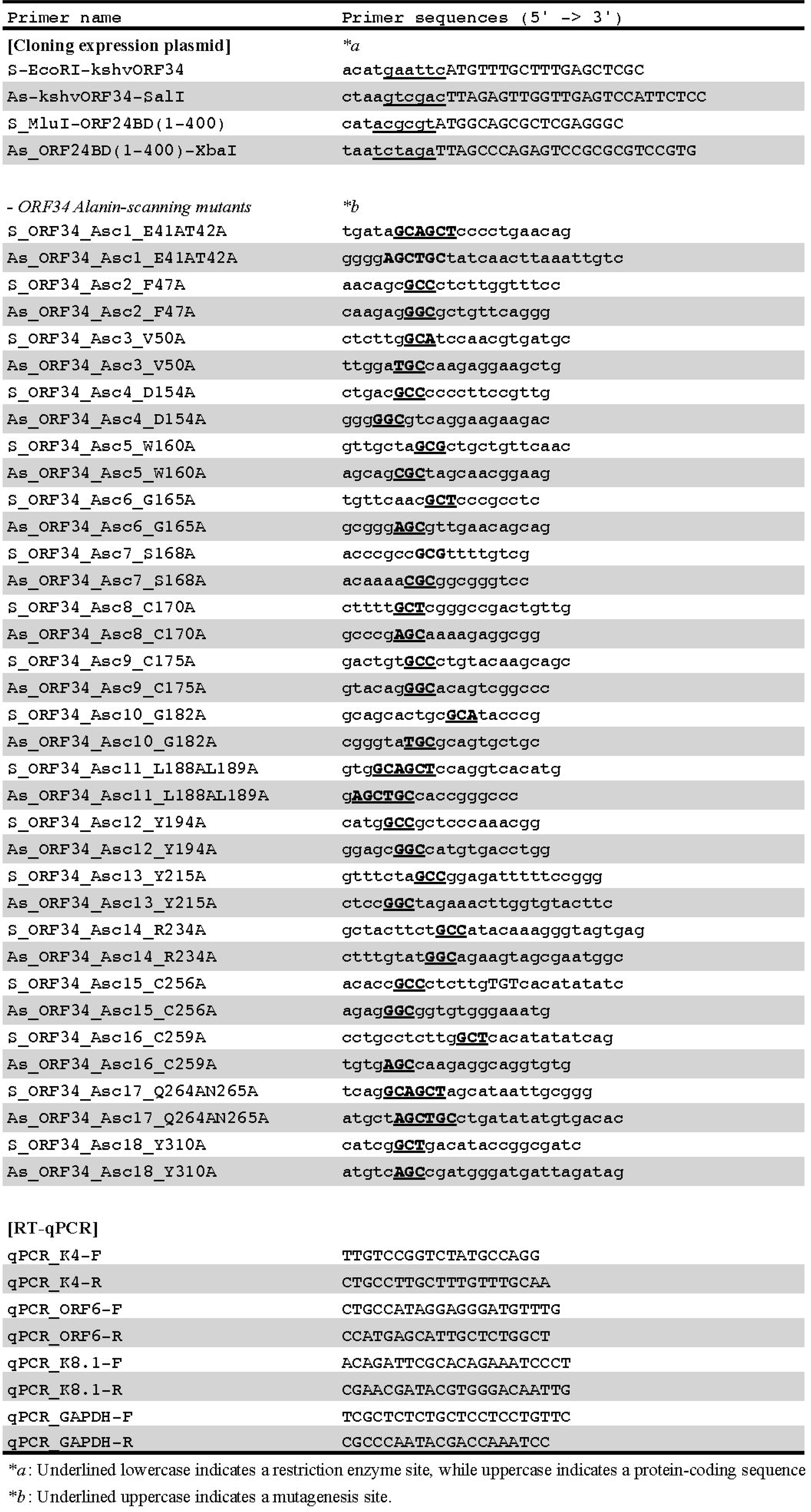
Primers for Construction of expression plasmids and RT-qPCR.

### Establishment of doxycycline-inducible recombinant KSHV-expressing cells and stably ORF34 WT- and mutant-expressing cells

For maintenance, iSLK cells were cultured in growth medium of DMEM/fetal calf serum 10% containing 1 μg/mL puromycin (Fujifilm-Wako Chemicals, Osaka, Japan) and 0.25 mg/mL of G418 (Fujifilm-Wako Chemicals). The KSHV BAC16 mutant (ΔORF34-BAC16) and its revertant (ΔORF34Rev-BAC16), as previously described (27), were transfected into iSLK cells using Screenfect A plus (Fujifilm-Wako Chemicals) according to the manufacturer’s instructions. Transfected cells were selected using 1000 μg/mL of hygromycin B (Fujifilm Wako Chemicals) to establish doxycycline-inducible recombinant KSHV-producing cell lines (iSLK-Δ34Rev and iSLK-Δ34).

To establish stable ORF34-expressing cells for complementation, pCI-blast-3xFLAG-ORF34 and empty vector pCI-blast-3xFLAG were transfected into iSLK-Δ34Rev and iSLK-Δ34 cells, and transfected cells were selected and maintained in 10 μg/mL and 7.5 μg/mL of Blasticidin S (Fujifilm-Wako Chemicals), respectively. Thus, stable cell lines iSLK-Δ34Rev/pCI-blast-3xFLAG, iSLK-Δ34/pCI-blast-3xFLAG, iSLK-Δ34/pCI-blast-3xFLAG-ORF34WT, and iSLK-Δ34/pCI-blast-3xFLAG-ORF34 mutants were established.

### Measurement of KSHV production

To quantify virus production, KSHV virions in the culture supernatant were quantified as described previously (27, 30, 50, 51) with minor modifications. Briefly, iSLK-harboring KSHV BAC cells were treated with 0.75 mM NaB and 4 μg/mL doxycycline for 72 h to induce lytic replication and production of recombinant KSHV, and culture supernatants were harvested. The culture supernatants (150 μL) were treated with DNase I (NEB, Ipswich, MA, U.S.A.) to obtain enveloped and encapsidated viral genomes. Viral DNA was purified and extracted from 100 μL of DNase I-treated culture supernatant using a QIAamp DNA Blood Mini Kit (QIAGEN GmbH, Hilden, Germany). To quantify viral DNA copies, SYBR Green real-time PCR was performed using THUNDERBIRD Next SYBR qPCR Mix (TOYOBO, Osaka, Japan) with KSHV-encoded ORF11 specific primers (27, 30).

### Quantification of viral gene expression

To assess the roles of ORF34 WT and its mutants in viral gene expression, RT-qPCR was performed as previously described (30). iSLK-Δ34 cells complemented with ORF34 WT or its mutants were stimulated and cultured, as described in the previous section. Total RNA was extracted from the treated cell lines using Sepasol-RNA II super (Nacalai Tesque, Inc.). Total RNA was digested with DNase using the ReverTra Ace qPCR RT Master Mix with gDNA Remover (TOYOBO) to exclude host and viral genomic DNA (gDNA), which interfered with mRNA quantification as a background. cDNA was synthesized using the same kit and subjected to SYBR green real-time PCR using the THUNDERBIRD Next SYBR qPCR Mix (TOYOBO) and the primer sets listed in Table 1, based on Fakhari and Dittmer (52) and the selected unique CDS referenced by Bruce *et al.* (53). Relative KSHV mRNA expression was determined using the GAPDH expression and ΔΔCt methods.

### Western blotting, pull-down assays, and antibodies

To detect viral proteins, iSLK-Δ34 cells complemented with ORF34 WT or mutants were stimulated and cultured, as described in the previous section. The treated cells were washed with PBS, directly lysed with W.B. sample buffer, and boiled. The W.B. samples were then subjected to western blotting, as described previously (30).

Pull-down assays between vPIC components and ORF34 were performed as previously described (27, 30, 35). Briefly, 293T cells were transfected with the indicated plasmids using PEI-MAX (54) and cultured for 2 days. Transfected cells were lysed using lysis buffer (20 mM HEPES (pH 7.9), 0.18 M NaCl, 0.1% (v/v) NP-40, 10% (v/v) Glycerol, 0.1 mM EDTA) with protease inhibitors and briefly sonicated to destroy the cell nucleus. Whole-cell extract, which is the supernatant obtained from the lysate centrifuged to discard cell debris, was subjected to affinity purification using S-protein-agarose (Merck Millipore, Burlington, MA, U.S.A.) for 1– 2 h and washed 4 times. Purified precipitates, including bait and prey proteins (2xS-tagged ORF34 WT/mutants and their counterparts) binding to S-protein-agarose, and whole-cell extracts were subjected to western blotting.

Treatment of the vPIC interaction with the zinc chelator TPEN (*N*,*N*,*N’*,*N’*-tetrakis (2-pyridylmethyl) ethylenediamine; TCI, Tokyo, Japan) was also performed as described previously (30). Briefly, 293T cell lysates transfected with 2xS-ORF34 or control vector were subjected to S-protein-agarose purification in the presence of each dose of TPEN or vehicle (ethanol). The beads associated with bait protein ORF34 WT or control were mixed with the cell lysate of prey proteins (6xMyc-ORF24BD or 3xFLAG-ORF66) and overexpressed in 293T cells in the presence of each dose of TPEN or vehicle. Beads were washed 4 times and subjected to western blot analysis.

Anti-Myc (9E10; Santa-Cruz, CA, U.S.A.), anti-S-tag pAb (MBL, Nagoya, Japan), anti-FLAG (DDDDK-tag) (FLA-1; MBL), Anti-HHV8 K-bZIP (F33P1; Santa-Cruz), Anti-HHV8 K8.1A/B (4A4; Santa Cruz), and anti-β-Tubulin (10G10; Fujifilm-Wako Chemicals) were used as the primary antibodies. HRP-linked anti-mouse IgG antibody (Jackson ImmunoResearch, Inc., West Grove, PA, U.S.A.) and HRP-linked anti-rabbit IgG antibody (Jackson ImmunoResearch, Inc.) were used as the secondary antibodies. Antibody-bound proteins were visualized using ECL Western Blotting Detection Reagents (ATTO, Tokyo, Japan) on an X-ray film (FUJIFILM Corp., Tokyo, Japan).

The pull-down efficiency score indicated that the band intensity of the prey protein (ORF24BD, ORF66) obtained from the mean region of interest using the ImageJ software program (Version; 2.0.0-rc-43/1.52n) was normalized to that of the bait protein (ORF34 WT/mutants).

### ChIP-qPCR

To survey the recruitment of ORF34 to the TSS of the L gene, iSLK-Δ34 cells complemented with ORF34 WT or mutants were subjected to ChIP with several minor modifications of previous methods (30). Complemented iSLK-Δ34 cells with ORF34 WT or mutants were stimulated and cultured as described above virus production section. A total of 1.2 × 10^6^ cells of each iSLK-Δ34-complemented cell line were seeded onto 10 cm dishes and stimulated with 10 mL of growth medium containing 0.75mM NaB and 4μG/mL Dox for 72 h. The cells were washed once and supplemented with 10 mL of pre-warmed 2% FCS medium. 11xFixation buffer (1 mL) containing methanol-free formaldehyde (#11850-14; Nacalai Tesque) (11.1% formaldehyde, 50 mM HEPES (pH8.0), 100 mM NaCl, 1 mM EDTA, 0.5 mM EGTA) was directly added to culture dishes and incubated for 10 min at room temperature, then sequentially quenched by 1mL of 1.5 M glycine. The fixed cells were washed with ice-cold PBS 3 times and Farnham Buffer (5mM PIPES (pH8.0), 85mM KCl, 0.5% (v/v) NP-40) with protease inhibitor cocktail (#25955-11; Nacalai Tesque) and scraped from dishes. The cells were homogenized in a 25-gauge syringe, and the pellet was washed with Farnham buffer. The cell nuclear pellets were suspended into 440 μL SDS-lysis buffer (50 mM Tris (pH8.0), 10 mM EDTA, 1% (w/v) SDS), sonicated by Bio-Raptor (High, 30 s; ON, 30 s; OFF, ×6 cycles) and centrifuged to remove debris. Cell nuclear extracts were diluted 10 times by ChIP Dilution Buffer (50 mM Tris (pH8.0), 167mM NaCl, 1.1% (v/v) Triton X-100, 0.11% (w/v) sodium deoxycholate). The diluted extracts were pre-cleared with 30 μL of Protein G Sepharose (Cytiva, Tokyo, Japan). The supernatants (1.2 mL) were subjected to immunoprecipitation with 30 μG of protein G magnetic beads (Dynabeads Protein G, Thermo Scientific) overnight. The magnetic beads were pre-conjugated with 2 mg/sample of normal mouse IgG (Santa Cruz Biotechnology) or anti-FLAG monoclonal antibody (M2; Sigma) for approximately 6-8 hours in 0.5% BSA/PBS solution. The IP-subjected magnetic beads were washed with RIPA buffer (50 mM Tris (pH8.0), 150 mM NaCl, 1mM EDTA, 1% (v/v) Triton X-100, 0.1% (w/v) SDS, 0.1% (w/v) sodium deoxycholate) twice, RIPA high-salt buffer (50 mM Tris (pH8.0), 500 mM NaCl, 1mM EDTA, 1% (v/v) Triton X-100, 0.1% (w/v) SDS, 0.1% (w/v) sodium deoxycholate), LiCl buffer (10 mM Tris (pH8.0), 250 mM LiCl, 1mM EDTA, 0.5% (v/v) NP-40, 0.5% (w/v) sodium deoxycholate), TE buffer (10 mM Tris (pH8.0), 1mM EDTA), and TE buffer containing 0.5% RIPA buffer. Finally, 120 μL of ChIP direct elution buffer (10 mM Tris (pH8.0), 300 mM NaCl, 5mM EDTA, 0.5% (w/v) SDS) was added to the magnetic beads and incubated overnight at 65°C for de-crosslinking. As a 1% input sample, 12 μL of the diluted extracts was diluted 10 times with ChIP direct elution buffer and subjected to de-crosslinking and after-treatments in parallel with the magnetic bead samples. The samples were digested with RNase A and Proteinase K at 37°C and 55°C, respectively. DNA was extracted from 110 μL of the digested supernatant using a Gel/PCR purification kit (Nihon Genetics) with minor modifications, column binding DNA was washed twice, and eluted with 75 μL of the kit elution buffer. The extracted DNA solutions were subjected to qPCR using primers for the K8.1-TSS (30).

### Statistical analyses and data visualization

The standard deviation was determined by analyzing the data obtained from 3 independent samples, and is indicated as error bars. Welch’s *t*-test was used to determine differences between the indicated groups. p-values are shown in the figure. For the functional profile analysis of ORF34 WT and its mutants, a heatmap with a dendrogram that highlights hierarchical clustering was visualized using R (Ver 4.0.3) and R Studio with R package “Superheat” (https://rlbarter.github.io/superheat/)(55).

## Data Availability

The data that support the findings of this study are available from the corresponding author, T.W., upon reasonable request.

## Acknowledgments

BAC16, a KSHV BAC clone, was a kind gift from Dr. Jae U Jung (Lerner Research Institute of Cleveland Clinic, U.S.A.). We also thank Dr. Gregory A. Smith (Northwestern University, U.S.A.) for providing the *E. coli* strain GS1783.

This work was partially supported by JSPS Grant-in-Aid for Scientific Research (C) (JP21K08509; T.W. and JP19K07669; S.O.), Takeda Science Foundation (T.W.), The Association for Research on Lactic Acid Bacteria (S.O.), the University of the Ryukyus Research Project Promotion Grant for Young Researchers (20SP04102; T.W.), the Research Start-up and the CAS Faculty Mellon Fund from American University (T.I.), and the National Institutes of Health through a grant from the National Institute of Allergy and Infectious Diseases (7R15AI172610; T.I.).

In this work, we would also like to thank Ms. Kazumi Tamaki, who is a member of the University of the Ryukyus integrated technology center, for her valuable technical support and documental management related to our laboratory work.

